# Long-term label-free assessments of individual bacteria using three-dimensional quantitative phase imaging and hydrogel-based immobilization

**DOI:** 10.1101/2022.06.04.494801

**Authors:** Jeongwon Shin, Jinho Park, Geon Kim, Moosung Lee, Yongkeun Park

**Affiliations:** Department of Biological Sciences, Korea Advanced Institute of Science and Technology (KAIST), Daejeon, 34141, South Korea; Department of Physics, KAIST, Daejeon, 34141, South Korea; KAIST Institute for Health Science and Technology, KAIST, Daejeon 34141, South Korea; Tomocube Inc., Daejeon 34051, South Korea

## Abstract

Three-dimensional (3D) quantitative phase imaging (QPI) enables long-term label-free tomographic imaging and quantitative analysis of live individual bacteria. However, the Brownian motion or motility of bacteria in a liquid medium produces motion artifacts during 3D measurements and hinders precise cell imaging and analysis. Meanwhile, existing cell immobilization methods produce noisy backgrounds and even alter cellular physiology. Here, we introduce a protocol that utilizes hydrogels for high-quality 3D QPI of live bacteria maintaining bacterial physiology. We demonstrate long-term high-resolution quantitative imaging and analysis of individual bacteria, including measuring the biophysical parameters of bacteria and responses to antibiotic treatments.

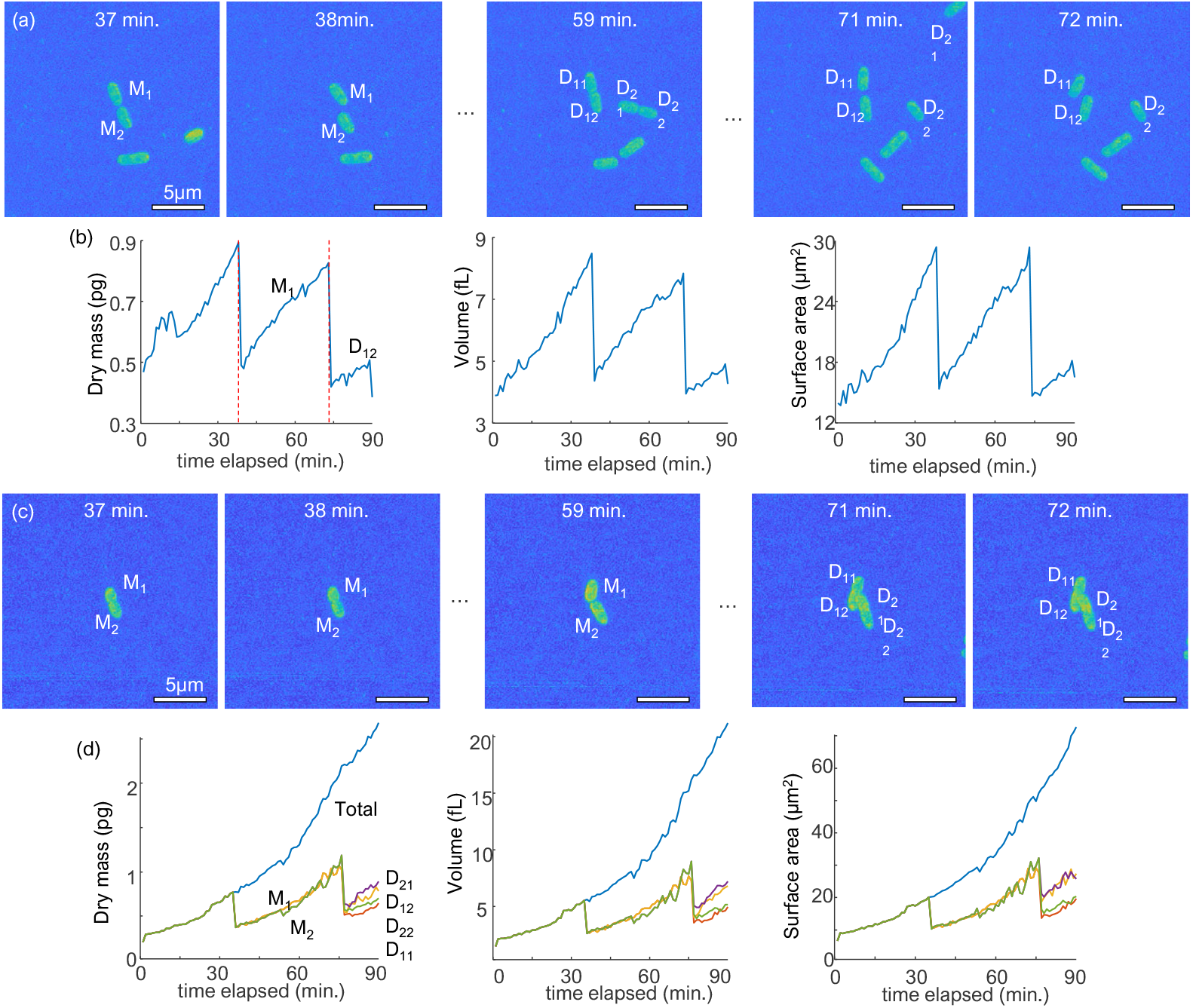

## 1. Introduction

Long-term monitoring of bacteria is a vital task in the healthcare, bioengineering industries, and biological sciences. In clinical settings, identifying pathogenic bacteria and determining effective antibiotics against bacteria are crucial processes in treating infectious diseases [1–3]. Moreover, engineering bacteria for the efficient production of biomaterials is an active field of research [4, 5]. Bacteria have also been utilized as model organisms to understand cellular mechanisms owing to their simplicity [6–8]. However, most conventional approaches to the study of bacteria have limited sensitivity because they rely on the bulk properties of bacterial colonies or high-concentration solutions. To identify pathogenic bacteria, time-consuming microbial culturing is required to secure sufficient signals in mass spectrometry-based detection devices [9]. Recently, advanced single-cell techniques have been developed and used to investigate individual bacteria [10–12]. Notably, the identification and analysis of bacteria at the single-cell level have been demonstrated in several studies [13–15].

Quantitative phase imaging (QPI) is a label-free imaging technique that has been exploited in various biological studies [16]. QPI enables label-free, high-contrast images by measuring the optical phase delay of scattered light [16]. By enabling investigations at the single-cell level, QPI has been utilized to study blood cells [17, 18], bacteria [19], cancer cells [20–22], SAR-VoC-2 virus [23], tissues [24, 25], and bacteria [13, 26]. Due to its high-resolution, label-free, quantitative imaging capability, 2D QPI techniques have been utilized for the study of bacteria. Bacteria are studied in more detail using three-dimensional (3D) QPI, which reconstructs the 3D refractive index (RI) tomogram from multiple 2D measurements of scattered light. 3D QPI facilitates single-cell level identification of bacteria [13], studying the response of antimicrobial photodynamic therapy [27], quantitative tracking of polymer synthesis in bacteria [28], and time-lapse investigation of antibiotic-treated bacteria [26].

However, the measurement of QPI images of individual bacteria is accompanied by experimental difficulties. One such difficulty is the Brownian motion and motility of bacteria in a liquid environment, which induce wobbling or traveling motion. The acquisition of high-quality 3D images is hindered in a liquid medium because bacterial motility generates imaging artifacts and prohibits long-term measurements [Fig. 1(a)]. Because 3D QPI reconstructs an RI tomogram from multiple 2D measurements, the image quality drastically drops with changes in the sample geometry during measurements. In addition, during long-term measurements, bacteria often travel from their original locations, occasionally out of the field of view (FOV).

**Fig. 1.**
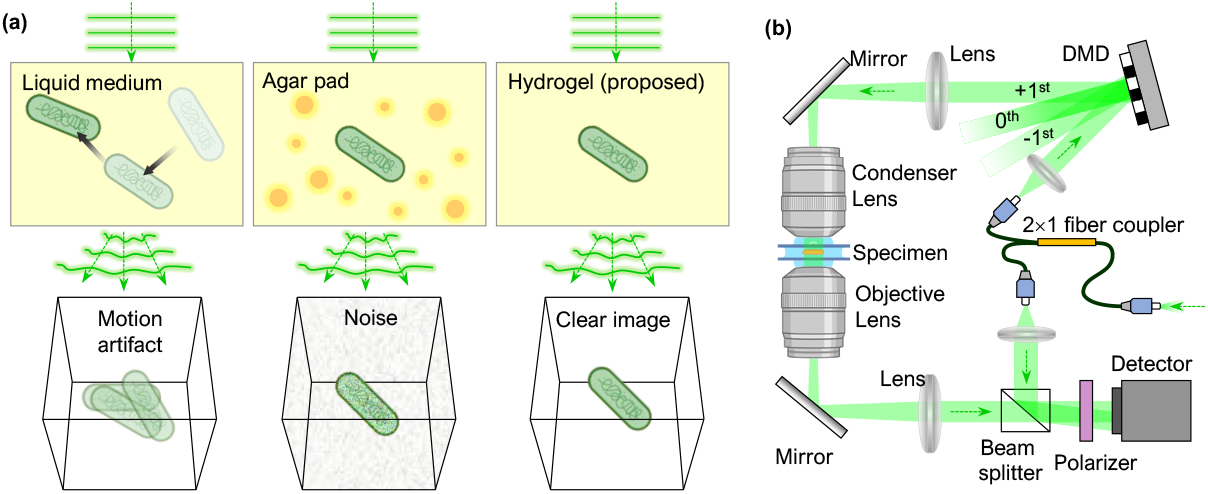
(a) QPI measurements of bacteria in different environments. The image quality is reduced by bacterial motility and background noise in a liquid medium and an agar pad, respectively. A hydrogel-based environment prevents both issues. (b) Schematic of the ODT system. DMD: digital micromirror device.

Several studies have suggested immobilization techniques to circumvent motility-related issues when measuring individual bacteria. One technique utilizes poly-L-lysine, which is frequently used to facilitate cell adhesion to plasticware or glass surfaces [29]. However, the physiological alteration of bacteria due to poly-L-lysine restricts the utility of this technique [30]. Another bacterial immobilization technique is an agar pad [31–33]. However, when applied to 3D QPI, an agar pad produces noisy images [Fig. 1] because of the inhomogeneous distribution of RI in an agar pad.

To achieve 3D QPI measurements of bacteria with neither motion-induced problems nor background noise, we propose an experimental method that utilizes a hydrogel to improve the imaging quality. Hydrogels are cross-linked hydrophilic polymers that form optically clear extracellular matrices and are suitable materials for addressing the aforementioned issues [34, 35]. Here, we experimentally demonstrate a method for imaging bacteria using hydrogel substrates and optical diffraction tomography (ODT), a 3D QPI technique [36]. Our method achieves higher image quality and stable sample immobilization compared to using a liquid medium or an agar pad environment. Moreover, our method facilitates long-term analyses of cellular features and investigation of the response to antibiotics. Based on the advantages emphasized in our study, our method provides a high-quality auxiliary technique for the study of individual bacteria.

## 2. Methods

### 2.1 Sample preparation

The bacterial strains *Klebsiella pneumoniae* (clinical isolates from the Asian Bacterial Bank of the Asia Pacific Foundation for Infectious Diseases) and *Escherichia coli* (ATCC25922) were stored in a −70°C deep freezer. Five microliters of the bacterial sample were added to 1000 μL of tryptic soy broth (TSB; MBcell) and incubated for 1 h at 37°C in a 5% CO_2_ shaking incubator. The stabilized mixture (100 μL) was spread on tryptic soy agar (TSA) plates and incubated for 24 h at 37°C in a 5% CO_2_ incubator. The TSA plates were stored at 4°C and used within a week. Before the experiments, a colony was selected, resuspended in 1000 μL TSB, and incubated for 24 h at 37°C in a 5% CO_2_ shaking incubator.

### 2.2 Hydrogel preparation

Hydrogels were formed by the combination of HyStem and Extralink (HYS020, Sigma-Aldrich, Saint Louis, Missouri, USA) with a volume ratio of 4:1. HyStem and Extralink are thiol-modified hyaluronan and thiol-reactive crosslinkers, respectively. HyStem and bacterial cultures were incubated in a 37°C water bath before use. For the 70% hydrogel sample, we mixed 14 μL of HyStem, 7.5 μL of bacteria culture, and 3.5 μL of Extralink in one Eppendorf (EP) tube and 14 μL of HyStem, 7.5 μL of TSB, and 3.5 μL of Extralink in another EP tube. For the 80% hydrogel sample, we mixed 16 μL of HyStem, 5 μL of bacterial culture, and 4 μL of Extralink in one EP tube and 16 μL of HyStem, 5 μL of TSB, and 4 μL of Extralink in another EP tube. For the 90% hydrogel sample, we mixed 18 μL of HyStem, 2.5 μL of bacterial culture, and 4.5 μL of Extralink in one EP tube and 18 μL of HyStem, 2.5 μL of TSB, and 4.5 μL of Extralink in another EP tube. We then placed a 20 mm × 20 mm cover glass on a TomoDish (Tomocube Inc., Daejeon, Korea), gently pressed only the sides of the glass to secure, and incubated the chamber on a 37°C hot plate. Ten microliters from each EP tube were added to each side groove of the TomoDish. Finally, the chamber was sealed by adding 10 μL of mineral oil to the grooves. TomoDish was incubated for 10 min on a hot plate at 37°C to fully solidify the hydrogel. HyStem and Extralink were stored at −20°C when not in use.

### 2.3 Agar pad preparation

Agar pads are agarose gels that solidify in thin cuboid forms. We prepared a 20 mm × 20 mm parafilm and excised a square hole of 12 mm × 12 mm. The parafilm was placed on a TomoDish and fixed by partial melting on a hot plate at 70°C. Fully melted 1.5% agarose gel (214010, BD, Franklin Lakes, New Jersey, USA) was poured onto the TomoDish and coverslipped. After the agarose gel was fully solidified, the coverslip was removed without generating any tears or wrinkles on the agar pad. Next, 5 μL of a sample was dropped on the agar pad, which was then gently covered with a new coverslip. Finally, the chamber was sealed by adding 10 μL of mineral oil to the grooves and incubated for 10 min on a 37°C hot plate.

### 2.4 RI diffraction tomography

To acquire a 3D RI tomogram of the bacteria, we used an ODT instrument (HT-2H, Tomocube Inc., Daejeon, Korea) with a live cell imaging chamber (Tomochamber, Tomocube Inc., Daejeon, South Korea). This system is based on a Mach-Zehnder interferometric microscope with a digital micromirror device (DMD) to control the angle of the sample beam [Fig. 1(b)] [37]. A coherent laser beam (*λ* = 532 nm in vacuum) was split into sample and reference beams using a 2×2 single-mode fiber optic coupler. The DMD controls the angle of the sample beam impinging on a sample. The beam diffracted by a sample is imaged on the camera plane, where it interferes with the reference beam. Using the phase-retrieval algorithm, the phase and amplitude images were retrieved from each interferogram. The 3D RI distribution was reconstructed from various 2D holograms of the sample from different illumination angles and by inversely solving the Helmholtz equation with the Rytov approximation. The theoretically calculated lateral and axial spatial resolutions of the optical imaging system were 110 nm and 360 nm, respectively. Because of the limited numerical aperture of condensers and objective lenses, side-scattering signals were not collected, leading to image quality degradation of the reconstructed 3D RI tomograms. To resolve this missing cone problem, an iterative regularization algorithm based on a non-negative constraint was used [38]. The detailed implementation and reconstruction code were as published previously [39, 40].

### 2.5 Cell doubling time analysis

Cell doubling time analysis was performed through manual inspection of the time-lapse maximum intensity projection (MIP) images. MIP is a projection method for 3D data that visualizes only the maximum RI values among RI from every parallel 3D data plane to the visualization plane. In each tomogram, the individual bacteria were automatically segmented using a threshold-based algorithm. The time point for each cell division was marked by manually inspecting the visualization of the segmentation masks. To secure highly precise statistical comparisons, for each bacterium we standardized the starting time point of the analysis. Specifically, the analysis started immediately after the first division was observed. In this way, variance in the initial cell phase between individual bacteria was excluded. We defined the timepoint of cell division as the time point when an additional segment of the cytoplasm occurred for the first time by division. This definition excludes the error of misrecognizing the time point when two daughter cells overlap and separate again in the MIP image. To avoid misrecognizing the division point if two daughter cells happened to overlap again, we used the interval between two adjacent cell division time points as the doubling time. For accuracy, measurements were performed three times independently. The analysis of cells in the liquid medium involved relatively immobilized cells.

### 2.6 Cellular features analysis

To measure cellular features, voxels with RI values higher than a set RI value were selected by masking the sample area. The dry mass of each cell was calculated using the linear relationship between RI and cellular dry mass [41, 42]: *n*(***r***) – *n*0 = *α* × *C*(***r***). Here *n, n*0, *α*, and *C* refer to the RI of a sample, the RI of the background medium, RI increment (RII), and protein concentration respectively. RII, which is an increment of an RI in a solution per an increment of concentration in a solute, determines the concentration of each point. The typical RII value for proteins is known to be 0.185 mL/g [43]. Since many bacteria have proteinaceous organelles [26], we utilized this typical RII value for proteins in our analysis. The cellular dry mass values were calculated by the total sum of the concentration of each voxel multiplied by its volume [26]. The cellular volume was estimated by the number of pixels and image resolution. Other physiological features were obtained in a similar manner. For the analyses of cells in the liquid medium, measurements were performed on the relatively immobilized cells.

## 3. Results

### 3.1 Measurements of high-quality ODT using a hydrogel environment

To validate the imaging capability of the hydrogel-based environment in 3D, we compared the 3D QPI images of individual bacteria obtained in a liquid solution, agar pad, and hydrogel [Fig. 2].

**Fig. 2.**
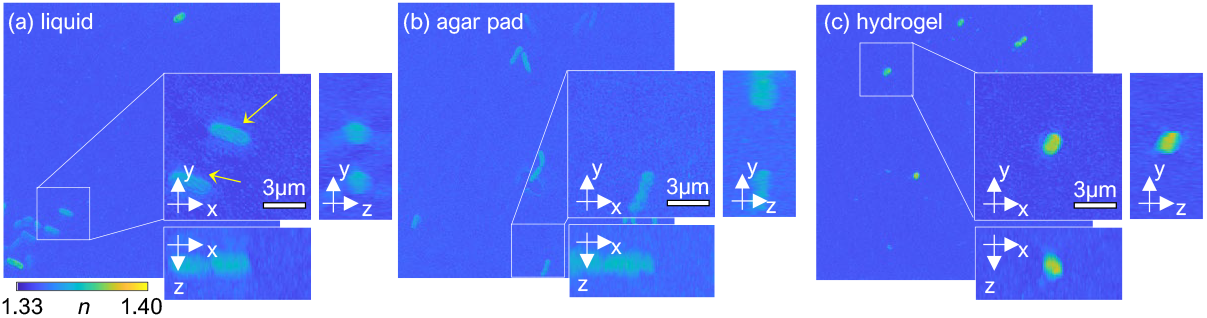
MIP visualization of 3D RI tomograms of *K. pneumoniae* in liquid (a), agar pad (b), and hydrogel (c).

The motion artifact in the hydrogel-based environment was dramatically reduced compared with that in the liquid environment. In liquid media, large numbers of bacteria exhibited wobbling motions during measurement producing motion artifacts; as a result, RI values were relatively underestimated, obscuring boundaries [Fig. 2(a), the yellow arrows]. The RI tomograms of bacteria in the agar pad showed speckle light background noise, implying inhomogeneous distribution of RIs in the agar pad [Fig. 2(b)]. In contrast, in the hydrogel environment, none of the bacteria were blurred [Fig. 2(c)]. Thus, our hydrogel-based protocol facilitates more accurate profiling of individual bacteria without the need for tedious data curation. For the remaining analyses in liquid media, we utilized curated tomograms without motion artifacts to address further issues, including background noise and motility.

We examined the background noise level of the 3D QPI image and the degree of bacteria immobilization in the liquid, agar pad, and hydrogel environment [Fig. 3]. 3D RI tomograms were measured for 1.5 hours in each medium environment and visualized using MIP with the corresponding bacterium centered in the image. *K. pneumoniae* in the liquid medium traveled and rotated [Fig. 3 (a)], whereas bacteria in the agar pad and the hydrogel were fixed, enabling monitoring of growth in different directions [Figs. 3(b) and 3(c)]. The hydrogel environment allows for reliable visualization and measurements of bacterial growth and division.

**Fig. 3.**
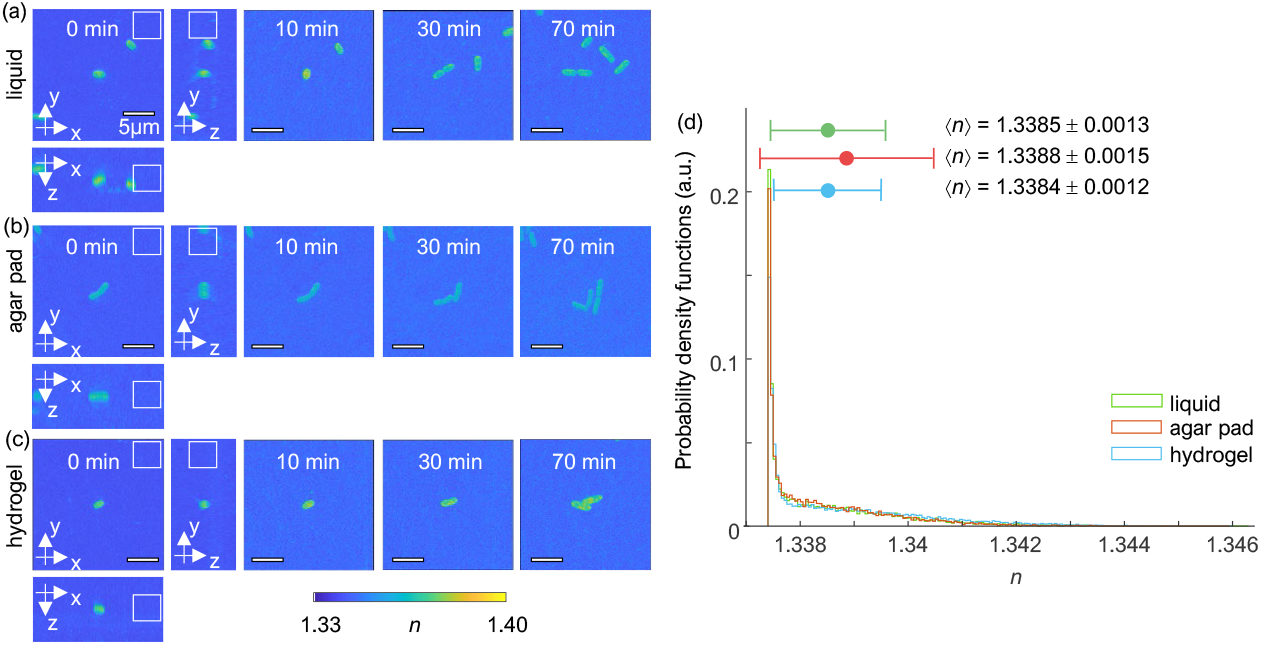
Time-lapse of MIP visualization of 3D RI tomograms of *K. pneumoniae* in different media: liquid medium (a), agar pad (b), and hydrogel (c). At 0 min, both 3D views of maximum RI projection and volumes marked with white squares which are used to calculate and visualize RI distribution of background as in (d) are indicated. (d) Probability density function plots of the distribution of background RIs measured from 3D RI tomograms at 0 min depending on media. For the background, we used cubes with a length of 40 pixels. Average and standard deviation in each plot are marked with circles and bars, respectively. For all cases, the lowest background RI values, which indicate pure media, were excluded to retain efficient data visualization. a.u., arbitrary unit.

Moreover, we analyzed the RI distribution of background values by measuring 3D RI tomograms at 0 min and generated probability density function plots [Fig. 3(d)]. As shown in Fig. 3(d), the distribution of background RIs for the hydrogel system assumed the lowest average and standard deviation compared to the other systems. This indicates low noise levels and evenness of the hydrogel system. By comparison, the agar pad system recorded the highest average and standard deviation of background RIs, demonstrating high noise levels. These results indicate that the hydrogel is an excellent system for bacteria immobilization while maintaining noise levels within the range of the liquid medium.

### 3.2 Biological compatibility of a hydrogel as a medium for bacteria culture

We then compared bacterial division in the hydrogel environment and in the liquid medium to assess the effect of hydrogel on the physiology of bacteria [Fig. 4]. No drastic physiological changes were observed, as the doubling time of bacteria in two environments did not vary significantly. In particular, we investigated cell morphology and the distribution of doubling time of *K. pneumoniae* in the liquid medium and in various concentrations of the hydrogel [Fig. 4]. We acquired RI tomograms at 1 min intervals and measured bacterial growth and division in the MIP images. Timepoints at division and at 3 min before and after division were chosen for the morphological analysis [Fig. 4(a)]. MIP images show that bacteria retain similar morphologies, grow to a fixed size, and reproduce through binary fission without any abnormal alterations in all conditions.

**Fig. 4.**
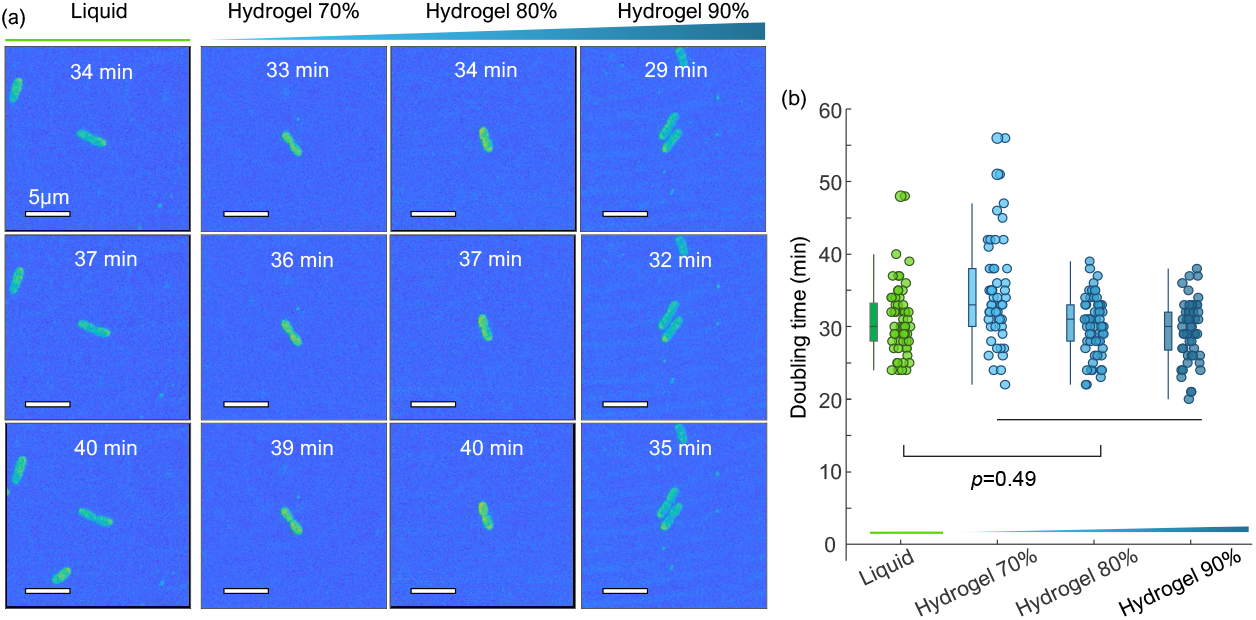
(a) MIP visualization of 3D RI tomograms of dividing *K. pneumoniae* over time in different media: a liquid medium, a hydrogel 70%, a hydrogel 80%, and a hydrogel 90% medium. (b) Distribution of *K. pneumoniae* doubling time in the different media. For the liquid and the pooled hydrogel environments, the result of the Wilcoxon rank-sum is given.

In a successive step, the doubling time of *K. pneumoniae* was obtained to quantitatively determine the effects of hydrogel on biological metabolism. The division events of 57, 50, 61, and 53 bacteria were investigated for each medium environment: liquid medium, hydrogel 70%, hydrogel 80%, and hydrogel 90% medium. To improve accuracy, we performed three separate examinations for each condition.

The distributions of doubling time of *K. pneumoniae* in the various media were comparable. The average and standard deviation of doubling time of *K. pneumoniae* in the liquid medium, hydrogel 70%, hydrogel 80%, and hydrogel 90% were 30.75 ± 4.58 min, 34.42 ± 6.95 min, 30.20 ± 3.98 min, and 29.53 ± 4.13 min, respectively [Fig. 4(b)]. The Wilcoxon rank-sum test *p*-values are 0.0025, 0.8142, and 0.3235 for 70%, 80%, and 90% hydrogel medium, respectively. The obtained *p*-value for the liquid and the pooled hydrogel media was 0.49, indicating statistical similarity. Thus, hydrogel, irrespective of polymer concentration, provides an environment similar to that of the liquid medium.

### 3.3 Cell tracking efficiency of hydrogel when yielding cellular features

Our hydrogel-based protocol facilitated the 1.5-hour monitoring of a single bacterium, allowing the investigation of morphological and biochemical features over time. The following cellular features were retrieved and analyzed using 3D RI tomograms: cellular dry mass, volume, and surface area. The temporal changes in these features were plotted to quantitatively assess bacterial growth in both the liquid and hydrogel media [Figs. 5(b) and 5(d)].

**Fig. 5.**
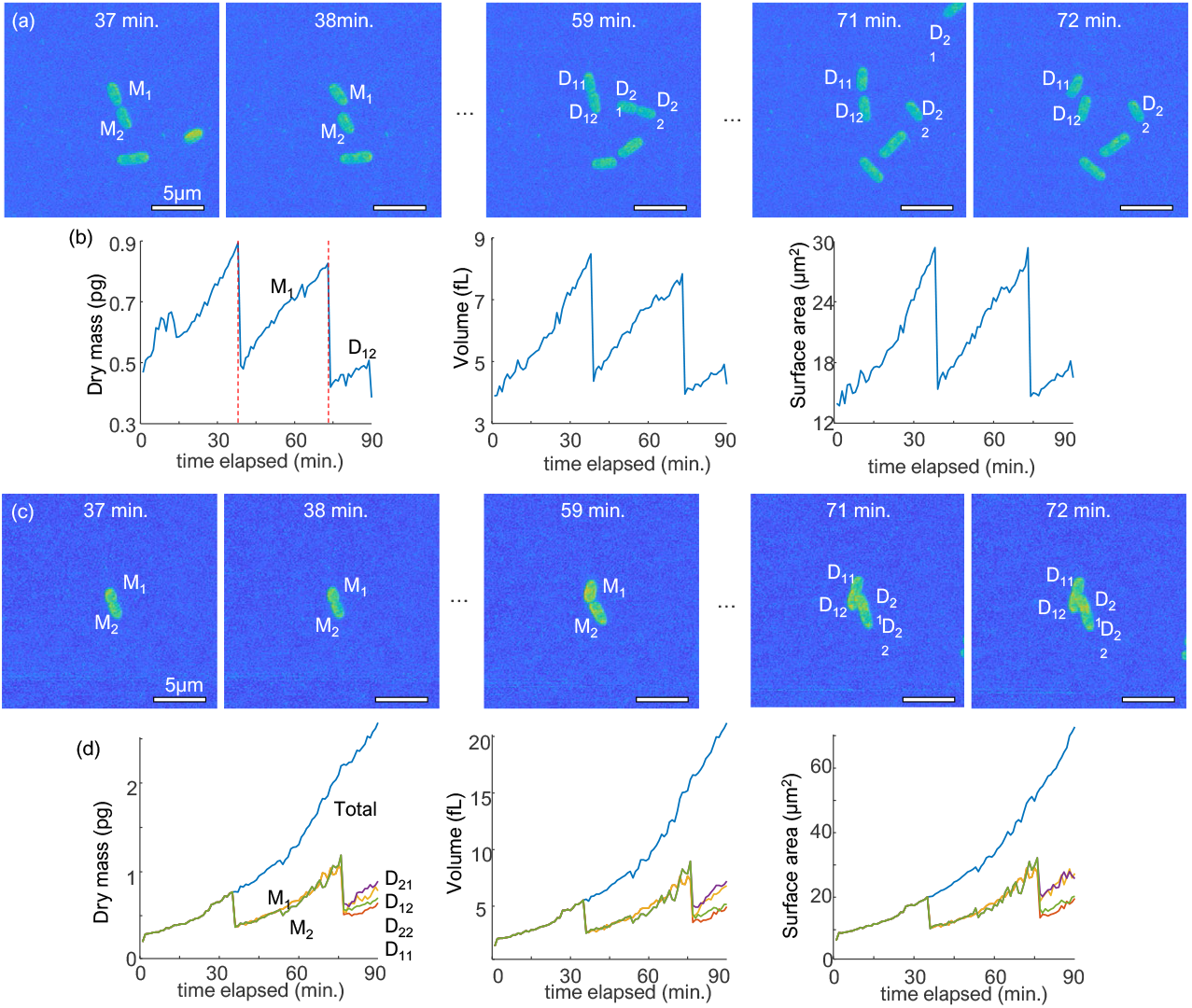
MIP visualization and quantitative analysis of 3D RI tomograms of *K. pneumoniae* in the liquid (a), (b) and hydrogel (c), (d) medium. The quantitative analysis includes cellular dry mass, volume, and surface area extracted from each snapshot tomogram. (a), (c) M1 and M2 refer to the mother cells while D11, D12 and D21, D22 refer to two daughter cells from each mother cell. In (a), one daughter cell (D21) can be seen at the upper right corner traveling out of the field of view at 71 min. (b) Time points showing sudden drops in cellular dry mass of *K. pneumoniae* are marked with red dashed lines. In (d), pooled data of all daughter cells and single-cell analyses are presented.

The growth of *K. pneumoniae* could be continuously tracked in the 80% hydrogel medium; this was difficult to accomplish in the liquid medium because of bacterial motility. The values of the cellular features in the liquid medium show sudden drops at certain points [Fig. 5(b)]. A review of acquired images showed two cases that hindered continuous growth measurements in the liquid medium. In the first case, the motility and rotation of the daughter cells rendered it difficult to trace their origins. In the second case, one of the daughter cells moved out of the field of view [Fig. 5(a), D21 at 71 min]. In contrast, the hydrogel medium becomes a practical environment for continuously tracking and analyzing cellular features. Bacteria are trackable and well immobilized for all daughter cells [Fig. 5(c)]. Thus, the pooled cellular features of the daughter cells in the hydrogel medium displayed a constant increase without sudden drops. In addition, analysis of the features of each daughter cell in the hydrogel gel was feasible [Fig. 5(d)]. The cellular features of single daughter cells clearly show two consecutive cell divisions, while analysis of the features of the cell clusters can potentially facilitate diagnosis.

### 3.4 Further application for imaging-based antimicrobial susceptibility testing

Finally, we demonstrated the potential application of a hydrogel as an imaging medium for imaging-based antimicrobial susceptibility testing (AST) [44, 45] (Fig. 6). We treated *K. pneumoniae* and *E. coli* with 200 μg/mL of ampicillin in the 80% hydrogel medium and measured 3D RI tomograms for 90 min to assess whether the hydrogel-based environment is suitable for evaluating bactericidal responses [26].

**Fig. 6.**
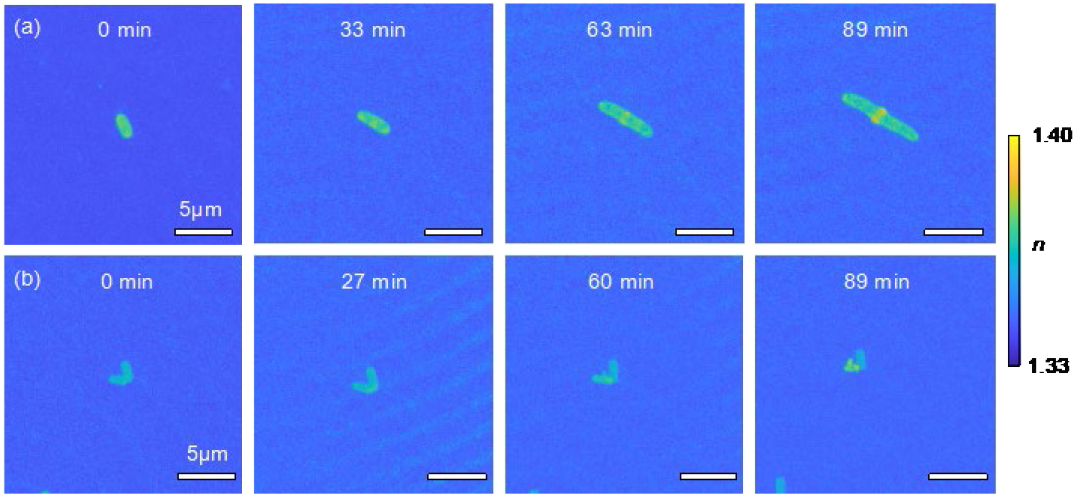
(a) MIP visualization of 3D RI tomograms of *K. pneumoniae* (a) and *E. coli* (b) treated with 200 μg/mL of ampicillin.

Several morphological alterations can be visualized. The MIP of *K. pneumoniae* treated with ampicillin shows that RI values decreased over time [Figure 6(a)]. Moreover, even after sufficient time, cell length increased without division [Fig. 6(a), 63 min]. In addition, the formation of a high RI bulge in the middle of the cell is evident [Fig. 6(a), 89 min]; this feature is the result of the inhibited transpeptidase [46, 47].

Figure 6(b) shows the MIP of a single *E. coli* treated with ampicillin. Over time, RI values decreased and a bulge formation can be detected [Fig. 6(b), 60 min] [46]. Based on these observations, we conclude that the hydrogel can be used successfully to image the physical responses of bacteria to antibiotics.

## 4. Discussion and Summary

We propose and demonstrate a protocol for 3D QPI of bacteria that does not suffer from motion-induced artifacts or background noise. Our protocol utilizes hydrogel as part of the loading medium and benefits from the formation of optically clear extracellular matrices [34, 35]. The hydrogel facilitates high-quality 3D QPI measurements of bacteria by immobilizing bacteria without generating noise. Moreover, the physiological characteristics of the samples remained constant, as indicated by our statistical analysis of the doubling time. The biocompatible and steady environment created by our protocol enabled long-term quantitative measurements. Further experiments also suggest the feasibility of our protocol in practical applications, such as measuring the efficacy of antibiotics in a hydrogel environment based on 3D QPI.

Identification of pathogenic bacteria is a potential application of the proposed method. Time-consuming microbial culture is an obstacle to early appropriate antibiotic treatment [48], and single-cell techniques, including QPI, aid in overcoming this limitation [13, 49]. Our method can aid in bacterial identification with QPI by providing a more stable environment for measurements. Our method can provide a high-quality imaging pipeline for the QPI-based identification of bacteria. An earlier study has demonstrated the rapid identification of bacteria using 3D QPI and deep learning [13], and we believe that our protocol can enhance the performance of such techniques by reducing motion artifacts and enabling long-term imaging. In addition, the classification of bacteria based on biomarkers will also be viable using our method [49], as it provides a more stable acquisition of biomarkers, including dry mass and volume. Therefore, our protocol will propel 3D QPI for rapid bacterial identification.

The study of bacterial responses in different environments is another field to which our protocol can contribute. One practical application of this method is AST. AST is used to assess the antibiotic resistance of specific bacteria. This information is crucial for medical doctors to determine the types of antibiotics used for infection treatment [50]. A commercialized label-free AST method evaluates samples in an agarose-based environment using bright-field microscopy. This method has the advantage of label-free imaging but still entails a day-long delay owing to the requirement of incubation [51, 52]. Using our approach, 3D QPI monitoring of individual bacteria can reveal antibiotic susceptibility within approximately an hour. Therefore, we envision that our protocol can accelerate label-free AST by eliminating the need for time-consuming cultivation procedures. Furthermore, a more efficient AST based on 3D QPI and our method can be realized by introducing microfluidic techniques. By forming drug gradients in a microfluidic chip, an on-chip AST, based on 3D QPI monitoring, can be facilitated.

Our method can also enhance the production of biopolymers by quantitatively monitoring the production levels of each cell. Bacteria can be used as cell factories for biopolymers, which are then used in medical and industrial applications [4]. By monitoring individual biopolymer production levels non-invasively, more efficient conditions for biopolymer production can be applied at medical and industrial levels [28]. Moreover, our protocol can aid in cellular mechanism studies of individual bacteria. Our method can facilitate investigating cellular physiology at single-bacterium resolution by providing long-term analysis of individual live cells [16, 53].

We plan to investigate the capability of our method extensively. Although we measured each specimen within 1.5 hours, with more precise maintenance of temperature and humidity, a much longer investigation would be possible. In addition, a hydrogel-based environment can be implemented in a shorter time using different polymer compositions, as already suggested by several studies [40, 41].

## References

1. J. O. Lay Jr, “MALDI-TOF mass spectrometry of bacteria,” Mass spectrometry reviews 20, 172–194 (2001).

2. S. Sauer, and M. Kliem, “Mass spectrometry tools for the classification and identification of bacteria,” Nature Reviews Microbiology 8, 74–82 (2010).

3. P. Seng, M. Drancourt, F. Gouriet, B. La Scola, P.-E. Fournier, J. M. Rolain, and D. Raoult, “Ongoing revolution in bacteriology: routine identification of bacteria by matrix-assisted laser desorption ionization time-of-flight mass spectrometry,” Clinical Infectious Diseases 49, 543–551 (2009).

4. M. F. Moradali, and B. H. Rehm, “Bacterial biopolymers: from pathogenesis to advanced materials,” Nature Reviews Microbiology 18, 195–210 (2020).

5. B. H. Rehm, “Bacterial polymers: biosynthesis, modifications and applications,” Nature Reviews Microbiology 8, 578–592 (2010).

6. S. Benzer, “On the topography of the genetic fine structure,” Proceedings of the National academy of Sciences of the United States of America 47, 403 (1961).

7. E. Russo, “Special Report: The birth of biotechnology,” Nature 421, 456–457 (2003).

8. F. R. Blattner, G. Plunkett, C. A. Bloch, N. T. Perna, V. Burland, M. Riley, J. Collado-Vides, J. D. Glasner, C. K. Rode, and G. F. Mayhew, “The complete genome sequence of Escherichia coli K-12,” science 277, 1453–1462 (1997).

9. E. Carbonnelle, C. Mesquita, E. Bille, N. Day, B. Dauphin, J.-L. Beretti, A. Ferroni, L. Gutmann, and X. Nassif, “MALDI-TOF mass spectrometry tools for bacterial identification in clinical microbiology laboratory,” Clinical biochemistry 44, 104–109 (2011).

10. P. C. Blainey, “The future is now: single-cell genomics of bacteria and archaea,” FEMS microbiology reviews 37, 407–427 (2013).

11. N. Musat, H. Halm, B. Winterholler, P. Hoppe, S. Peduzzi, F. Hillion, F. Horreard, R. Amann, B. B. Jørgensen, and M. M. Kuypers, “A single-cell view on the ecophysiology of anaerobic phototrophic bacteria,” Proceedings of the National Academy of Sciences 105, 17861–17866 (2008).

12. Z. Long, E. Nugent, A. Javer, P. Cicuta, B. Sclavi, M. C. Lagomarsino, and K. D. Dorfman, “Microfluidic chemostat for measuring single cell dynamics in bacteria,” Lab on a Chip 13, 947–954 (2013).

13. G. Kim, D. Ahn, M. Kang, Y. Jo, D. Ryu, H. Kim, J. Song, J. S. Ryu, G. Choi, and H. J. Chung, “Rapid and label-free identification of individual bacterial pathogens exploiting three-dimensional quantitative phase imaging and deep learning,” BioRxiv, 596486 (2019).

14. S. Pahlow, S. Meisel, D. Cialla-May, K. Weber, P. Rösch, and J. Popp, “Isolation and identification of bacteria by means of Raman spectroscopy,” Advanced drug delivery reviews 89, 105–120 (2015).

15. W. E. Huang, R. I. Griffiths, I. P. Thompson, M. J. Bailey, and A. S. Whiteley, “Raman microscopic analysis of single microbial cells,” Analytical chemistry 76, 4452–4458 (2004).

16. Y. Park, C. Depeursinge, and G. Popescu, “Quantitative phase imaging in biomedicine,” Nature photonics 12, 578–589 (2018).

17. G. Kim, Y. Jo, H. Cho, H.-s. Min, and Y. Park, “Learning-based screening of hematologic disorders using quantitative phase imaging of individual red blood cells,” Biosensors and Bioelectronics 123, 69–76 (2019).

18. J. Yoon, Y. Jo, M.-h. Kim, K. Kim, S. Lee, S.-J. Kang, and Y. Park, “Identification of non-activated lymphocytes using three-dimensional refractive index tomography and machine learning,” Scientific reports 7, 1–10 (2017).

19. M. Mir, Z. Wang, Z. Shen, M. Bednarz, R. Bashir, I. Golding, S. G. Prasanth, and G. Popescu, “Optical measurement of cycle-dependent cell growth,” Proceedings of the National Academy of Sciences 108, 13124–13129 (2011).

20. T. H. Nguyen, S. Sridharan, V. Macias, A. Kajdacsy-Balla, J. Melamed, M. N. Do, and G. Popescu, “Automatic Gleason grading of prostate cancer using quantitative phase imaging and machine learning,” Journal of biomedical optics 22, 036015 (2017).

21. V. K. Lam, T. C. Nguyen, V. Bui, B. M. Chung, L.-C. Chang, G. Nehmetallah, and C. B. Raub, “Quantitative scoring of epithelial and mesenchymal qualities of cancer cells using machine learning and quantitative phase imaging,” Journal of biomedical optics 25, 026002 (2020).

22. L. Kastl, M. Isbach, D. Dirksen, J. Schnekenburger, and B. Kemper, “Quantitative phase imaging for cell culture quality control,” Cytometry Part A 91, 470–481 (2017).

23. N. Goswami, Y. R. He, Y.-H. Deng, C. Oh, N. Sobh, E. Valera, R. Bashir, N. Ismail, H. Kong, and T. H. Nguyen, “Label-free SARS-CoV-2 detection and classification using phase imaging with computational specificity,” Light: Science & Applications 10, 1–12 (2021).

24. A. Bokemeyer, P. R. Tepasse, L. Quill, P. Lenz, E. Rijcken, M. Vieth, N. Ding, S. Ketelhut, F. Rieder, and B. Kemper, “Quantitative phase imaging using digital holographic microscopy reliably assesses morphology and reflects elastic properties of fibrotic intestinal tissue,” Scientific reports 9, 1–11 (2019).

25. H. Hugonnet, Y. W. Kim, M. Lee, S. Shin, R. H. Hruban, S.-M. Hong, and Y. Park, “Multiscale label-free volumetric holographic histopathology of thick-tissue slides with subcellular resolution,” Advanced Photonics 3, 026004 (2021).

26. J. Oh, J. S. Ryu, M. Lee, J. Jung, S. Han, H. J. Chung, and Y. Park, “Three-dimensional label-free observation of individual bacteria upon antibiotic treatment using optical diffraction tomography,” Biomedical optics express 11, 1257–1267 (2020).

27. A. Pietrowska, I. Hołowacz, A. Ulatowska-Jarża, M. Guźniczak, A. K. Matczuk, A. Wieliczko, M. Wolf-Baca, and I. Buzalewicz, “The Enhancement of Antimicrobial Photodynamic Therapy of Escherichia Coli by a Functionalized Combination of Photosensitizers: In Vitro Examination of Single Cells by Quantitative Phase Imaging,” International Journal of Molecular Sciences 23, 6137 (2022).

28. S. Y. Choi, J. Oh, J. Jung, Y. Park, and S. Y. Lee, “Three-dimensional label-free visualization and quantification of polyhydroxyalkanoates in individual bacterial cell in its native state,” Proceedings of the National Academy of Sciences 118 (2021).

29. G. E. Fantner, R. J. Barbero, D. S. Gray, and A. M. Belcher, “Kinetics of antimicrobial peptide activity measured on individual bacterial cells using high-speed atomic force microscopy,” Nature nanotechnology 5, 280–285 (2010).

30. K. Colville, N. Tompkins, A. D. Rutenberg, and M. H. Jericho, “Effects of poly (L-lysine) substrates on attached Escherichia coli bacteria,” Langmuir 26, 2639–2644 (2010).

31. G. Joyce, B. D. Robertson, and K. J. Williams, “A modified agar pad method for mycobacterial live-cell imaging,” BMC Research Notes 4, 1–4 (2011).

32. J. C. Locke, and M. B. Elowitz, “Using movies to analyse gene circuit dynamics in single cells,” Nature Reviews Microbiology 7, 383–392 (2009).

33. J. R. Moffitt, J. B. Lee, and P. Cluzel, “The single-cell chemostat: an agarose-based, microfluidic device for high-throughput, single-cell studies of bacteria and bacterial communities,” Lab on a Chip 12, 1487–1494 (2012).

34. M. T. Spang, and K. L. Christman, “Extracellular matrix hydrogel therapies: in vivo applications and development,” Acta biomaterialia 68, 1–14 (2018).

35. R. Singh, K. Shitiz, and A. Singh, “Immobilization of Bacterial Cells in Hydrogels Prepared by Gamma Irradiation for Bioremoval of Strontium Ions,” Water, Air, & Soil Pollution 231, 1–10 (2020).

36. E. Wolf, “Three-dimensional structure determination of semi-transparent objects from holographic data,” Optics communications 1, 153–156 (1969).

37. S. Shin, K. Kim, J. Yoon, and Y. Park, “Active illumination using a digital micromirror device for quantitative phase imaging,” Optics letters 40, 5407–5410 (2015).

38. J. Lim, K. Lee, K. H. Jin, S. Shin, S. Lee, Y. Park, and J. C. Ye, “Comparative study of iterative reconstruction algorithms for missing cone problems in optical diffraction tomography,” Optics express 23, 16933–16948 (2015).

39. S. Shin, K. Kim, T. Kim, J. Yoon, K. Hong, J. Park, and Y. Park, “Optical diffraction tomography using a digital micromirror device for stable measurements of 4D refractive index tomography of cells,” in Quantitative Phase Imaging //(International Society for Optics and Photonics 2016), p. 971814.

40. K. Kim, H. Yoon, M. Diez-Silva, M. Dao, R. R. Dasari, and Y. Park, “High-resolution three-dimensional imaging of red blood cells parasitized by Plasmodium falciparum and in situ hemozoin crystals using optical diffraction tomography,” Journal of biomedical optics 19, 011005 (2013).

41. R. Barer, “Determination of dry mass, thickness, solid and water concentration in living cells,” Nature 172, 1097–1098 (1953).

42. G. Popescu, Y. Park, N. Lue, C. Best-Popescu, L. Deflores, R. R. Dasari, M. S. Feld, and K. Badizadegan, “Optical imaging of cell mass and growth dynamics,” American Journal of Physiology-Cell Physiology 295, C538–C544 (2008).

43. C. Park, S. Shin, and Y. Park, “Generalized quantification of three-dimensional resolution in optical diffraction tomography using the projection of maximal spatial bandwidths,” JOSA A 35, 1891–1898 (2018).

44. R. Datar, S. Orenga, R. Pogorelcnik, O. Rochas, P. J. Simner, and A. van Belkum, “Recent advances in rapid antimicrobial susceptibility testing,” Clinical chemistry 68, 91–98 (2022).

45. A. van Belkum, C.-A. D. Burnham, J. W. Rossen, F. Mallard, O. Rochas, and W. M. Dunne, “Innovative and rapid antimicrobial susceptibility testing systems,” Nature Reviews Microbiology 18, 299–311 (2020).

46. B. G. Spratt, “Distinct penicillin binding proteins involved in the division, elongation, and shape of Escherichia coli K12,” Proceedings of the National Academy of Sciences 72, 2999–3003 (1975).

47. E. Efstratiou, A. I. Hussain, P. S. Nigam, J. E. Moore, M. A. Ayub, and J. R. Rao, “Antimicrobial activity of Calendula officinalis petal extracts against fungi, as well as Gram-negative and Gram-positive clinical pathogens,” Complementary Therapies in Clinical Practice 18, 173–176 (2012).

48. M. D. Gonsalves, and Y. Sakr, “Early identification of sepsis,” Current infectious disease reports 12, 329–335 (2010).

49. Y. Jo, J. Jung, J. W. Lee, D. Shin, H. Park, K. T. Nam, J.-H. Park, and Y. Park, “Angle-resolved light scattering of individual rod-shaped bacteria based on Fourier transform light scattering,” Scientific reports 4, 1–6 (2014).

50. Z. A. Khan, M. F. Siddiqui, and S. Park, “Current and emerging methods of antibiotic susceptibility testing,” Diagnostics 9, 49 (2019).

51. J. Choi, Y.-G. Jung, J. Kim, S. Kim, Y. Jung, H. Na, and S. Kwon, “Rapid antibiotic susceptibility testing by tracking single cell growth in a microfluidic agarose channel system,” Lab on a Chip 13, 280–287 (2013).

52. J.-H. Kim, T. S. Kim, H. G. Jung, C. K. Kang, K.-I. Jun, S. Han, D. Y. Kim, S. Kwon, K.-H. Song, and P. G. Choe, “Prospective evaluation of a rapid antimicrobial susceptibility test (QMAC-dRAST) for selecting optimal targeted antibiotics in positive blood culture,” Journal of Antimicrobial Chemotherapy 74, 2255–2260 (2019).

53. R. Kasprowicz, R. Suman, and P. O’Toole, “Characterising live cell behaviour: Traditional label-free and quantitative phase imaging approaches,” The international journal of biochemistry & cell biology 84, 89–95 (2017).

